# Alleviation of neuropathic pain by over-expressing a soluble colony-stimulating factor 1 receptor

**DOI:** 10.1101/2020.09.09.288928

**Authors:** Svetlana Gushchina, Ping K. Yip, Glesni A. Parry, Haripriya Sivakumar, Jie Li, Min Liu, Xuenong Bo

**Affiliations:** Centre for Neuroscience, Surgery and Trauma, Blizard Institute, Barts and the London School of Medicine and Dentistry, Queen Mary University of London, London E1 2AT, UK.; Department of Cytology, Histology and Embryology, Ogarev Mordovia State University, Saransk 430005, Republic of Mordovia, Russia.; Department of Laboratory Medicine, Taihe Hospital, Hubei University of Medicine, Shiyan, China.

**Keywords:** adeno-associated viral vector, colony stimulating factor-1 receptor, macrophage /microglia, neuropathic pain

## Abstract

In this study, we aim to alleviate neuropathic pain by suppressing microgliosis and macrophage accumulation, which is achieved by over-expressing a non-functional soluble colony-stimulating factor 1 receptor using adeno-associated virus 9 vectors (AAV9/sCSF1R). AAV9/sCSF1R and AAV9/GFP were intrathecally administered into mouse lumbar spine. Two weeks later, these mice underwent partial sciatic nerve ligation to induce neuropathic pain. GFP and sCSF1R were highly expressed in dorsal root ganglia (DRG) and spinal cords in AAV9-injected mice. Nerve ligation alone or pre-treated with AAV9/GFP led to significant microgliosis in the lumbar spinal cords and macrophage accumulation in DRG and sciatic nerves. In nerve-ligated mice pre-treated with AAV9/sCSF1R the microglia densities in the dorsal and ventral horns and macrophage densities in DRG and sciatic nerves were significantly lower compared to nerve-ligated mice pre-treated with AAV9/GFP. Behavioural tests showed that nerve-ligated mice pre-treated with AAV9/sCSF1R had a significantly higher paw withdrawal threshold, indicating the alleviation of neuropathic pain. The results implicate that viral vector-mediated expression of sCSF1R may represent a novel strategy in long-term alleviation of neuropathic pain.

## Introduction

Millions of people worldwide suffer from debilitating chronic neuropathic pain, which significantly affects the quality of life of the patients. Neuropathic pain is caused by a lesion or a disease of the somatosensory nervous system such as peripheral nerve injury, diabetic neuropathy, spinal cord injury, multiple sclerosis, or herpes zoster virus infection (Bouhassira & Attal, 2019). The underlying mechanisms for neuropathic pain have been studied extensively for many years - however, many patients do not respond effectively to current treatment (Cohen & Mao, 2014). It was observed many years ago that peripheral nerve injury can produce rapid and profound mechanical allodynia, and proliferation and activation of microglia and astrocytes in the corresponding segments of the spinal cord (Colburn *et al*, 1999; Winkelstein *et al*, 2001). Furthermore, proliferation and activation of microglia were observed in other pain models such as diabetic pain and bone cancer pain (Tsuda *et al*, 2008; Yang *et al*, 2015). These findings indicate that proliferation and activation of microglia contribute towards the development of neuropathic pain after peripheral nerve injury (Inoue & Tsuda, 2018). Microgliosis after peripheral nerve injury was reported to be required for the maintenance of chronic neuropathic pain as well (Echeverry *et al*, 2017). An early study indicates that certain signalling molecule(s) released from injured neurons in dorsal root ganglia (DRG) into the dorsal horn of spinal cord induces microgliosis (Colburn *et al*., 1999). It was only recently that the signalling molecule was identified to be colony-stimulating factor 1 (CSF1) (Gu *et al*, 2016; Okubo *et al*, 2016). Ligation or partial transection of mouse sciatic nerve branches can lead to a significant increase of CSF1 expression in DRG (Guan *et al*, 2016). The CSF1 mRNA level was increased in DRG neurons while the CSF1 receptor (CSF1R) mRNA level in microglia in the spinal cord was increased in a rat model of partial sciatic nerve injury (Okubo *et al*., 2016). Such findings indicate that blocking CSF1 signalling may be an effective approach in treating nerve injury-induced neuropathic pain. Indeed, it was reported that CSF1R inhibitors were effective in alleviating neuropathic pain by blocking the proliferation of microglia in the dorsal horn in a partial sciatic nerve ligation model (Lee *et al*, 2018). However, the clinical use of CSF1R inhibitors for treating neuropathic pain may result in severe adverse effects, as CSF1 signalling is crucial for the physiological proliferation and functions of monocyte-derived cells such as macrophages and dendritic cells, and osteoclasts (Stanley *et al*, 1997). Moreover, apart from microgliosis in the spinal cord, macrophages infiltrated into DRG were also reported to contribute to neuropathic pain induced by peripheral nerve injury (Yu *et al*, 2020). It is, therefore, imperative to develop an alternative approach to target the CSF1 signalling pathway in the nervous system to treat neuropathic pain and other neurological disorders in which microglia play a role in the pathophysiology (Pons & Rivest, 2018).

CSF1R comprises an extracellular CSF1 binding domain, a transmembrane domain, and an intracellular C-terminal with a tyrosine kinase domain. To neutralize CSF1 released in the DRG and spinal cord, we over-expressed a soluble form of CSF1R (sCSF1R) which consists of only the extracellular domain with the CSF1 binding site. We postulate that this non-functional sCSF1R over-expressed via an adeno-associated viral vector 9 (AAV9) in the DRG and spinal cord would act as a decoy for CSF1 and compete with its binding to native CSF1R on microglia and macrophages, therefore, reducing the proliferation of microglia and macrophages due to the increased level of CSF1.

In this study, we derived a molecular therapy with a three-pronged approach by preventing the proliferation of microglia in the spinal cord, and the accumulation of macrophages in DRG and the injured nerves in mice with neuropathic pain induced by partial sciatic nerve ligation.

With this approach, we aim to achieve long-term pain alleviation without significantly affecting the microglia in other parts of the CNS, and the peripheral macrophages and dendritic cells.

## Results

### sCSF1R is secreted efficiently from transfected HEK293 cells

A molecular construct expressing sCSF1R with the nerve growth factor signal peptide fused to the N-terminal and HA-2A-GFP sequence at the C-terminal was generated (Fig 1A). The expression and secretion of the sCSF1R-HA-2A-GFP construct were initially tested on HEK293 cells, with the transfected cells immunostained with both anti-CSF1R and anti-HA tag antibodies. GFP fluorescence was visualized directly and mainly located in the nuclei (Fig 1B and C) while sCSF1R immunoreactivity (-ir) (Fig 1B) and HA-ir (Fig 1C) was predominantly located in the Golgi apparatus and secretory vesicles (yellow arrows in Fig 1C), indicating that sCSF1R and GFP molecules were separated during translation. Moreover, this was confirmed by immunoblotting the conditioned medium of sCSF1R-transfected cells. Detection with anti-GFP antibody showed a band of about 115 kDa (green arrowhead in the first lane in Fig 1D), the predicted molecular mass of the glycosylated sCSF1R-HA-2A-GFP fusion protein. Since the anti-GFP antibody used has a very high affinity, the band of sCSF1R-HA-2A-GFP appeared much stronger than those detected with anti-CSF1R and anti-HA antibodies (green arrowheads in Lane 2 and 3 in Fig 1D). In the lanes detected with anti-CSF1R and anti-HA antibodies, the main bands corresponded to the molecular mass of sCSF1R-HA-2A (orange arrowheads in Lane 2 and 3 of Fig 2D), indicating that most of the sCSF1R-HA-2A molecules were separated from GFP and secreted into the medium while most GFP molecules were retained inside the cells. The presence of sCSF1R molecules in the conditioned medium was further confirmed by immunoprecipitation. sCSF1R was immunoprecipitated using anti-HA antibody and detected with anti-CSF1R antibody (Lane 4 in Fig 1D). Secreted sCSF1R maintains its capability to bind to CSF1, as it was shown that the recombinant murine CSF1 (mCSF1) could be co-immunoprecipitated with sCSF1R in the conditioned medium using anti-HA antibody and detected with anti-mCSF1 antibody (Fig 1E).

**Figure 1.**
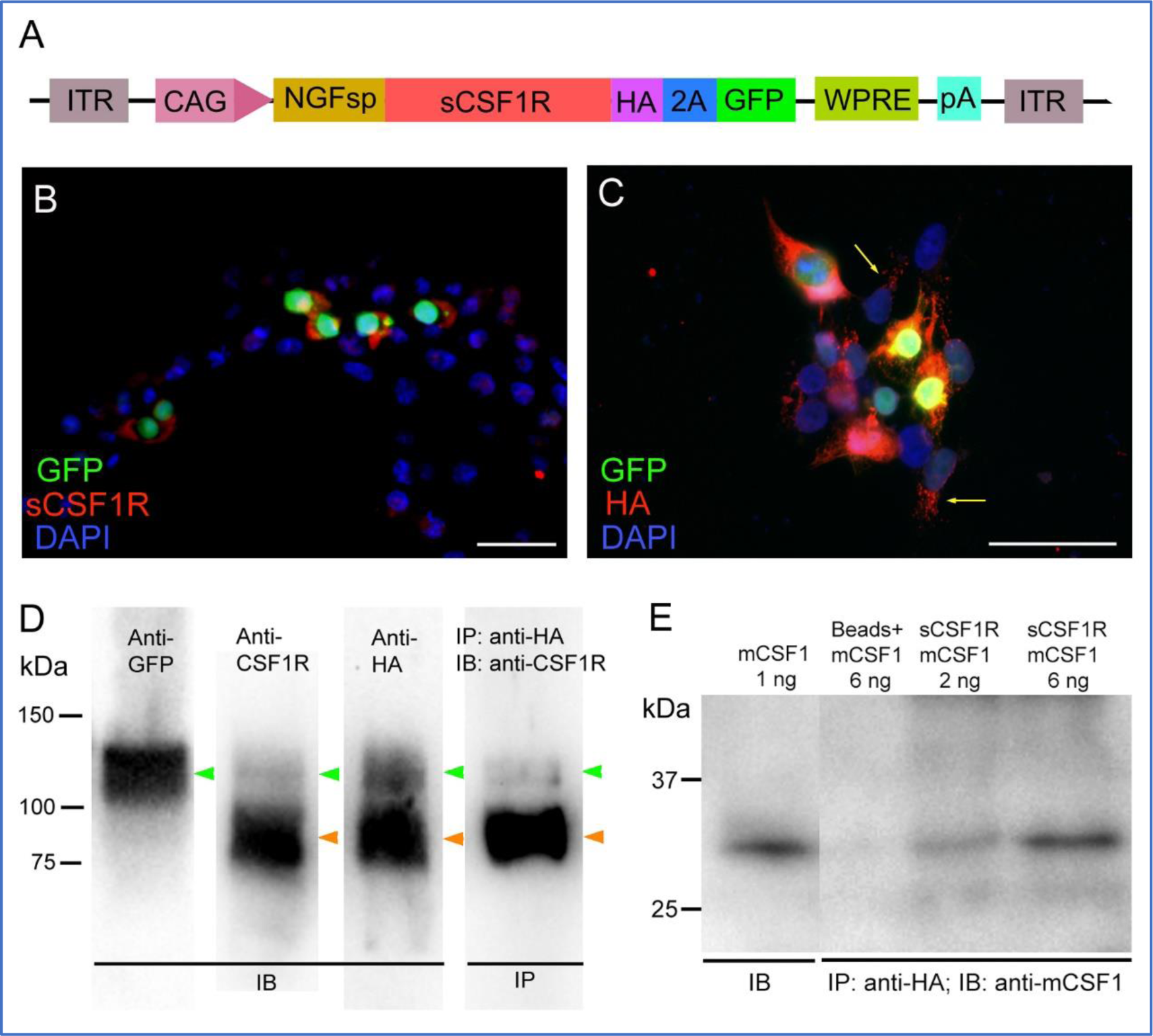
sCSF1R was secreted efficiently from transfected HEK293 cells. A Schematic diagram of AAV construct for sCSF1R. ITR: inverted terminal repeat of AAV; CAG: hybrid chicken β-actin promoter; NGFsp: signal peptide of nerve growth factor; HA: hemagglutinin epitope tag; 2A: self-cleaving peptide; WPRE: woodchuck hepatitis post-transcriptional regulatory element; pA: polyA. B, C HEK293 cells transfected with pAAV/sCSF1R-HA-2A-GFP and immunostained with anti-CSF1R antibody (B) or anti-HA (C). Yellow arrows in (C) point to sCSF1R in secretory vesicles in HEK293 cells. D Immunoblot showing sCSF1R in conditioned medium from sCSF1R transfected HEK293 cells detected by anti-GFP, anti-CSF1R, and anti-HA antibodies. Lane 4 shows sCSF1R immunoprecipitated using anti-HA antibody and detected with anti-CSF1R antibody. Green arrowheads in Lane 1 - 4 point to the un-cleaved sCSF1R-HA-2A-GFP fusion protein. Orange arrowheads in Lane 2 - 4 point to sCSF1R-HA-2A with GFP cleaved. E Immunoblot showing recombinant murine CSF1 (mCSF1) co-immunoprecipitated with sCSF1R in the conditioned medium using anti-HA antibody and detected with anti-mCSF1 antibody. Lane 1 was loaded with 1 ng mCSF1 as control. IB: immunoblotting; IP: immunoprecipitation. Data information: Scale bar, 25 µm in (B) and (C).

**Figure 2.**
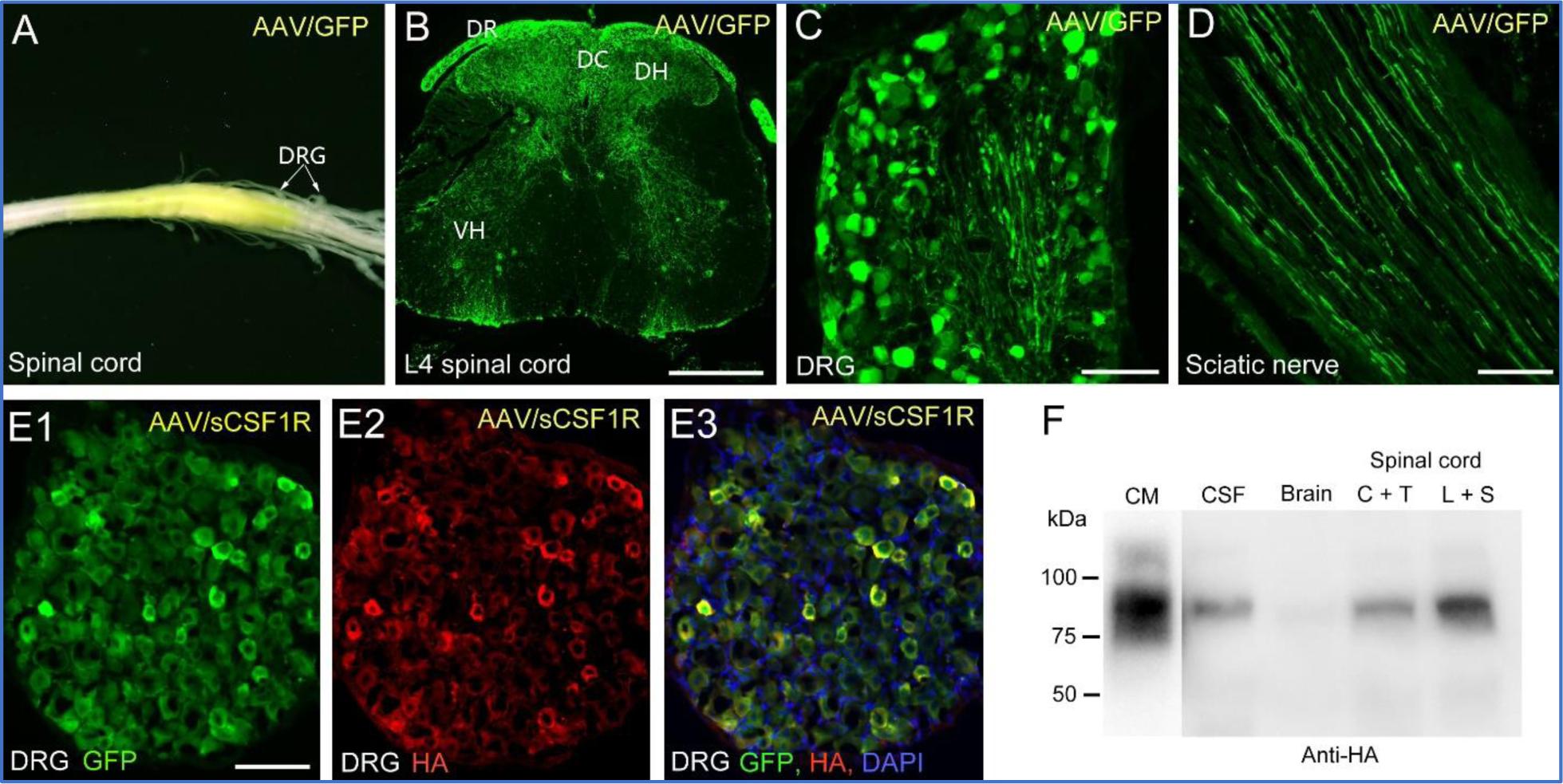
Expression of GFP and sCSF1R in the nervous system after intrathecal injection of AAV9/GFP or AAV9/sCSF1R. A A spinal cord with AAV9/GFP transduction viewed under non-artificial UV light showing high-level expression of GFP in the lumbar and sacral spinal cord and the associated DRG. B A cross-section of L4 spinal cord of AAV9/GFP-injected mouse showing dense GFP^+^ nerve fibres in the dorsal horn (DR), dorsal column (DC), dorsal roots (DR), and other regions of the spinal cord. Some neuronal bodies were GFP^+^ as well. C High-level expression of GFP was seen in neurons of all sizes in an L4 DRG of an AAV9/GFP-injected mouse. D A longitudinal section of the sciatic nerve from an AAV9/GFP injected mouse showing many GFP^+^ axons. E Injection of AAV9/sCSF1R led to expression of GFP (E1) and HA-tagged sCSF1R (E2) in DRG. Merged image (E3) shows that GFP and HA-tagged sCSF1R were co-expressed in the same cells. F Immunoblot of sCSF1R detected with an anti-HA tag antibody shows a higher level of sCSF1R in the lumbar and sacral spinal cord tissue than in the cervical and thoracic spinal cord tissue, cerebrospinal fluid, and brain tissue. Conditioned medium (CM) from sCSF1-GFP transfected HEK293 cells was used as control. Data information: Scale bar, 400 μm in (B); 100 µm in (C-E).

### AAV9 mediates high-level expression of sCSF1R in DRG and spinal cord

For *in vivo* studies, AAV9 vectors expressing the molecular construct NGFsp-sCSF1R-HA-2A-GFP (AAV9/sCSF1R) and GFP molecule only as control (AAV9/GFP) were produced. To test the efficiency of AAV9 mediated expression of sCSF1R and GFP in DRG and the spinal cord, the two viral vectors were intrathecally injected into the lumbar spines of mice. Two weeks after intrathecal injection of AAV9/GFP and AAV9 sCSF1R, the expression of GFP and sCSF1R in DRG, spinal cord, brain, and sciatic nerve were examined. It was found that AAV9 mediated a very high-level expression of the two transgenes. GFP expression in the spinal cord and the associated DRG of an AAV9/GFP injected mouse could be visualized directly without an artificial UV light source (Fig 2A) and the expression of GFP was mainly in the lumbar and sacral spinal cord, indicating that the injected AAV9 vectors predominantly transduced the cells close to the injection site. A cross-section of a lumbar spinal cord shows that GFP^+^ nerve fibres were concentrated in the dorsal horn and dorsal column (Fig 2B), indicating that most of them were the central branches of DRG neurons. GFP^+^ sensory axons also projected to other regions of the spinal cord including the ventral horn (Fig 2B, Fig EV1A). Some neurons in the dorsal and ventral horns were transduced, albeit the number of transduced cells was low, indicating that intrathecally injected AAV9 vectors at the dorsal surface of the spinal cord were not very efficient in penetrating deep into the spinal cord. Interestingly, GFP^+^ ascending sensory axons were observed in all sensory tracts in the thoracic and cervical spinal cord, medulla, and cerebellum (Fig EV1B-D). Also, some GFP^+^ neurons were observed in various regions of the brain, indicating some AAV9 vectors spread to the brain.

DRG neurons of all sizes were transduced by AAV9/GFP with very high efficiency (Fig 2C). Many GFP^+^ axons were seen in the sciatic nerve (Fig 2D). In DRG of AAV9/sCSF1R injected mice, many DRG neurons were also transduced, as demonstrated by positive immunostaining using anti-GFP and anti-HA tag antibodies (Fig 2E). Although GFP-ir overlapped HA-ir in the cytoplasm, more GFP-ir was seen in the nuclei.

As most of the neurons transduced by AAV9/sCSF1R after intrathecal injection were DRG neurons, we anticipate that sCSF1R would be released into the spinal cord where many central branches of DRG neurons terminate. Furthermore, any release of molecules by the DRG would be present in the surrounding cerebrospinal fluid. Indeed, immunoblotting of the supernatants of the spinal cord and brain homogenates showed that the levels of sCSF1R correlated to the densities of GFP^+^ axons in those regions, with the lumbar and sacral spinal cord containing the highest level of sCSF1R, while the brain had the lowest level (Fig 2F). Also, a moderate level of sCSF1R was detected in the cerebrospinal fluid.

### Over-expression of sCSF1R suppresses CSF1 expression in DRG neurons and spinal motor neurons after partial sciatic nerve ligation

To generate neuropathic pain, a commonly used peripheral nerve injury model involving the partial ligation of the sciatic nerves was carried out (Seltzer *et al*, 1990). We first examined the expression of CSF1 protein in the lumbar DRG and spinal cord. In normal mice, the levels of CSF1 in the DRG and spinal cord were rather low (Fig 3A and H), similar to what we have observed in the spinal cords in normal mice previously reported (Gushchina *et al*, 2018). Furthermore, CSF1 is secreted upon synthesis and diffuses into intracellular space, making it difficult to detect CSF1 inside the cells using immunohistochemistry. Sham surgery did not induce a significant increase in CSF1 expression in DRG and the spinal cord (Fig 3B and I). However, partial sciatic nerve ligation did lead to an increased CSF1 expression at various degrees in individual DRG neurons (Fig 3C) and motor neurons in the dorsal horn ipsilateral to the injury (Fig 3J). In sham-operated mice with the AAV9/GFP injection increased expression of CSF1 was observed only in a few DRG neurons (Fig 3D), but not in motor neurons in the ventral horn (Fig 3K). In nerve-ligated mice with AAV9/GFP injection increased CSF1 expression was observed in some DRG neurons (Fig 3E) and motor neurons in the ventral horn (Fig 3L). In contrast, in sham-operated mice with the AAV9/sCSF1R injection, no neuron with a detectable level of CSF1 expression was observed in DRG (Fig 3F) and the spinal cord (Fig 3M).

**Figure 3.**
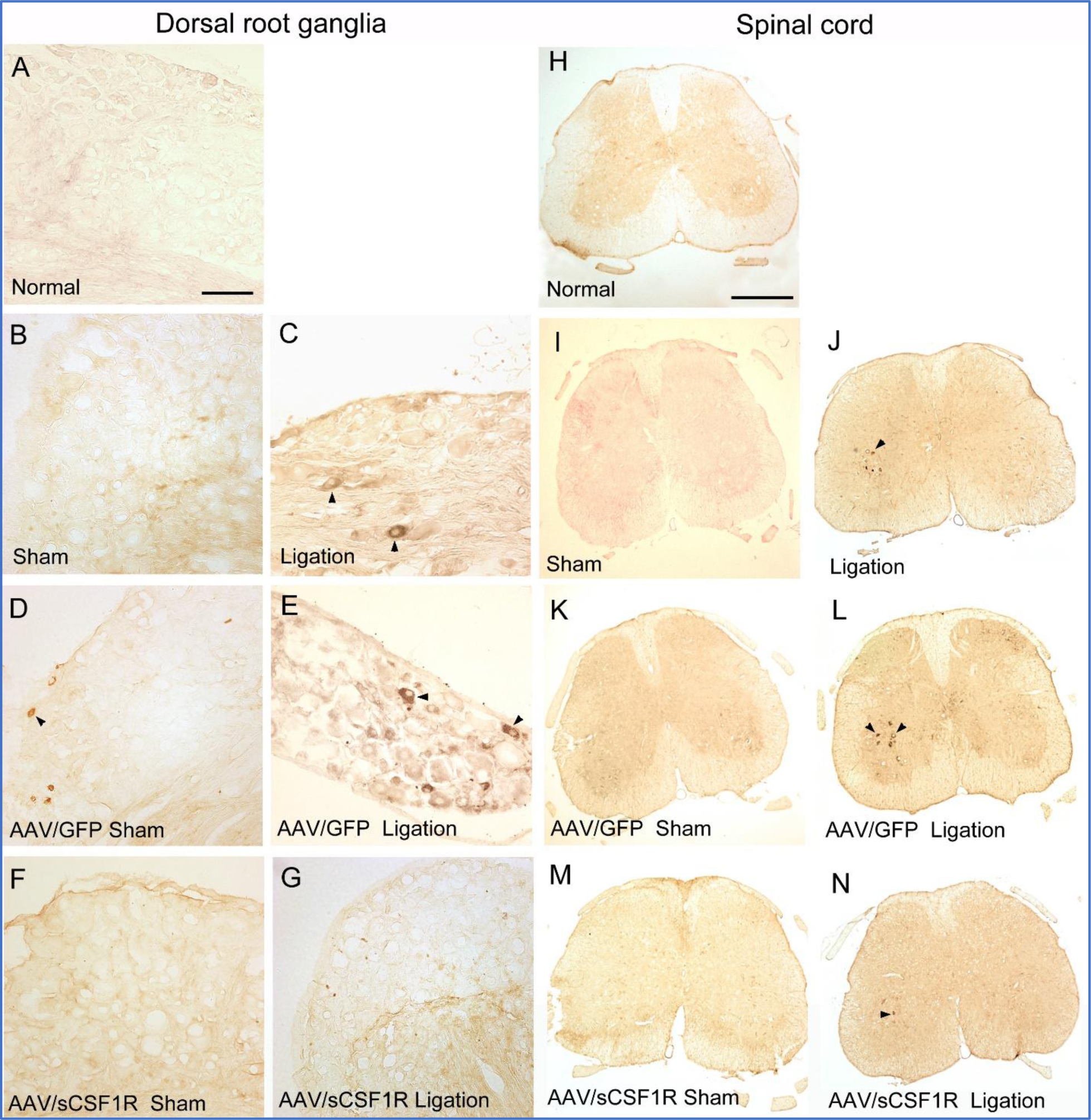
Over-expression of sCSF1R suppressed CSF1 expression in DRG neurons and motor neurons in the spinal cord after partial sciatic nerve ligation. Expression of CSF1 in DRG: A Normal mouse. B Mouse with sham surgery. C Mouse with partial sciatic nerve ligation. D Mouse received AAV9/GFP injection and sham surgery. E Mouse received AAV9/GFP injection and partial sciatic nerve operation. F Mouse received AAV9/sCSF1R injection and sham surgery. G Mouse received AAV9/sCSF1R injection and partial sciatic nerve ligation. Expression of CSF1 in the lumbar spinal cord: H Normal mouse. I Mouse with sham surgery. J Mouse with partial sciatic nerve ligation. K Mouse received AAV9/GFP injection and sham surgery. L Mouse received AAV9/GFP injection and partial sciatic nerve operation. M Mouse received AAV9/sCSF1R injection and sham surgery. N Mouse received AAV9/GFP injection and partial sciatic nerve ligation. Data information: arrowheads point to some neurons with high-level CSF1. Scale bar, 100 µm in (A-G); 400 µm in (H-N).

Furthermore, in nerve-ligated mice with AAV9/sCSF1R injection, there was no obvious increase in CSF1 expression in DRG (Fig 3F) and only a slight increase of CSF1 in a few motor neurons in the ventral horn (Fig 3N). These results were surprising as we did not anticipate that over-expression of sCSF1R in DRG neurons would affect the expression of CSF1 in DRG and motor neurons.

### Over-expression of sCSF1R reduces microgliosis in the spinal cord induced by partial sciatic nerve ligation

Microglia in the cross-sections of lumbar spinal cords were labelled with an anti-Iba1 antibody. The integrated densities that represent the areas and intensities of Iba1-ir were quantitated. Sham surgery did not significantly increase the microglia densities when compared with the normal spinal cord (Fig 4A, B and H). In contrast, partial sciatic nerve ligation induced significant microgliosis in the dorsal and ventral horns on the ipsilateral side (Fig 4C and H). In the ventral horn, the increased density of microglia was mainly in the area within the location of large motor neurons that showed significantly increased CSF1 expression after nerve ligation (Fig 3J). In AAV9/GFP-injected mice, sham surgery did not increase the microglia density significantly (Fig 4D and H), while nerve ligation significantly increased microglia density in the ipsilateral dorsal and ventral horns (Fig 4E and H). Interestingly microgliosis was also seen in the dorsal horns on the contralateral side (Fig 4E, Fig EV2A-D), but not significantly in the ventral horn. In AAV9/sCSF1R-injected mice, sham surgery did not increase the microglia density significantly (Fig 4F and H) and the microglia densities in both dorsal and ventral horns were significantly lower than those in AAV9/GFP-injected mice with sham surgery. In the spinal cord of the AAV9/sCSF1R-injected mice with nerve ligation, microglia densities in the dorsal and ventral horns were significantly lower than those of the AAV9/GFP-injected mice with nerve ligation, although they were still higher than those of the AAV9/sCSF1R injected mice with sham surgery (Fig 4G and H). These results indicate that over-expression of sCSF1R effectively reduced the microgliosis in the spinal cord, although the suppression is not complete at the viral dose used in this experiment.

**Figure 4.**
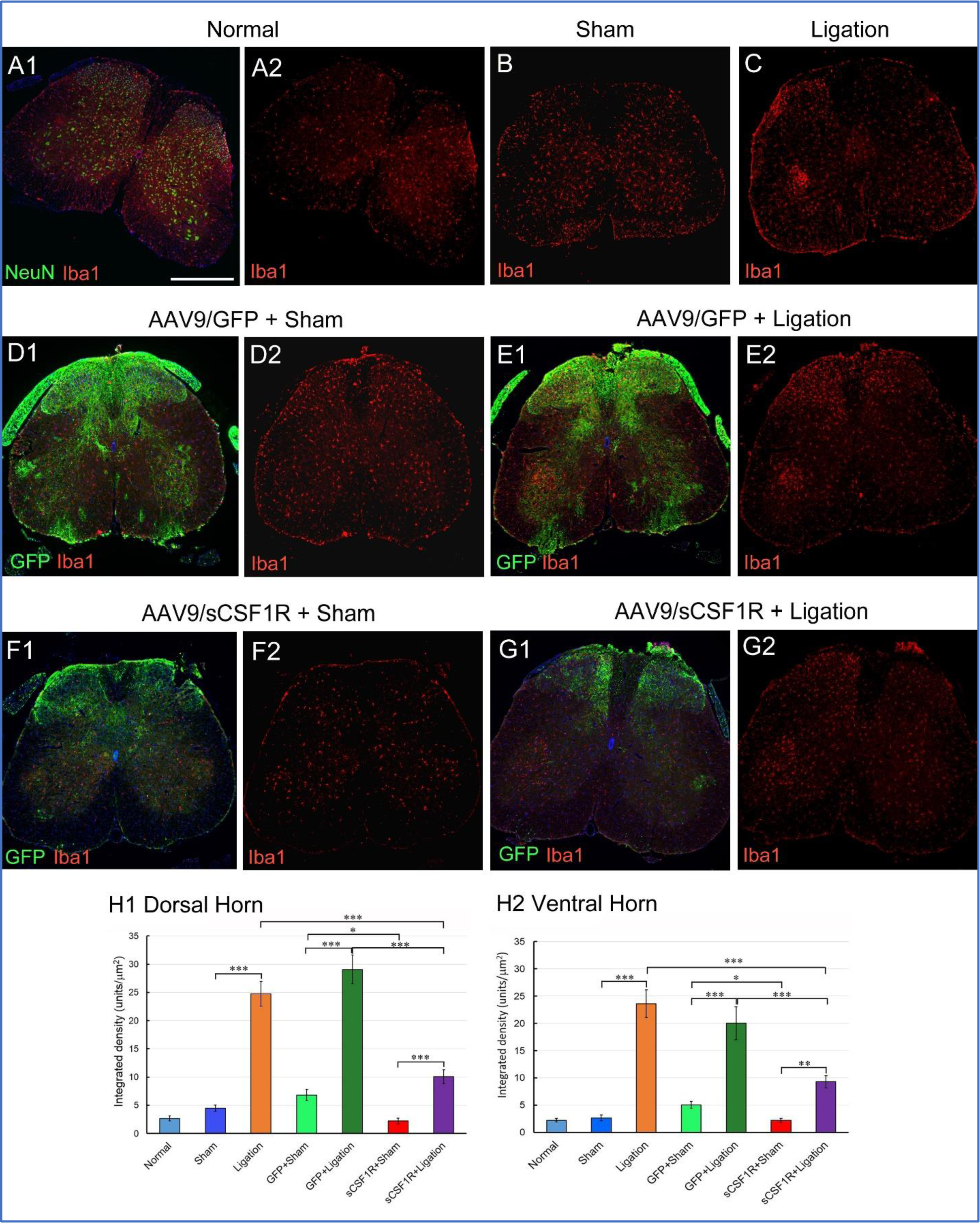
Over-expression of sCSF1R suppressed microgliosis in the spinal cord after partial sciatic nerve ligation. Photomicrographs shown in this figure were taken from the L4 spinal cord. 1, merge images; 2, red-channel only images. A1-A2 Normal B Sham surgery C Partial sciatic nerve ligation. D1-D2 AAV9/GFP + sham surgery. E1-E2 AAV9/GFP + ligation. F1-F2 AAV9/sCSF1R + sham surgery. G1-G2 AAV9/sCSF1R + ligation. H1-H2 Quantification of intensities of Iba1 immunoreactivity in the dorsal and ventral horns on the ipsilateral sides of the spinal cords. n = 6 per group. Data information: Scale bar, 400 μm in (A1), also applies to the other photomicrographs. Data are presented as mean ± SEM. \**p* < 0.05; \*\**p* < 0.01; \*\*\**p* < 0.001. Statistic analysis was performed using one-way ANOVA with post doc Tukey test.

### Over-expression of sCSF1R suppresses macrophage accumulation in DRG after partial sciatic nerve ligation

Macrophages in DRG were also labelled using anti-Iba1 antibody and the integrated densities of Iba1-ir were measured. A small number of Iba1^+^ macrophages were present in the normal DRG (Fig 5A). Sham surgery was able to induce a moderate but significant increase in the density of Iba1^+^ macrophages (Fig 5B and H), while partial sciatic nerve ligation induced significantly more macrophages than sham surgery (Fig 5C and H). Sham-operated mice with AAV9/GFP injection had an increased density of macrophages (Fig 5D and H), but not significantly higher than the sham surgery-only group. Nerve-ligated mice with AAV9/GFP injection showed a very significant increase in the density of macrophages (Fig 5E and H, Fig EV3E-H). Sham-operated mice with AAV9/sCSF1R injection expressed similar macrophage density to the normal and the sham surgery groups (Fig 5F and H), but significantly lower than that of the sham-operated mice with AAV9/GFP injection, indicating that sCSF1R was capable of reducing the sham surgery-induced slight increase in macrophage density in DRG. Nerve-ligated mice with AAV9/sCSF1R injection exhibited a moderately increased macrophage density when compared with the sham-operated mice with AAV9/sCSF1R injection, but a significantly lower density when compared with the nerve-ligated mice with AAV9/GFP injection (Fig 5G and H). These results indicate that over-expression of sCSF1R can significantly suppress the accumulation of macrophages in DRG, leading to a reduced inflammatory response to partial nerve ligation.

**Figure 5.**
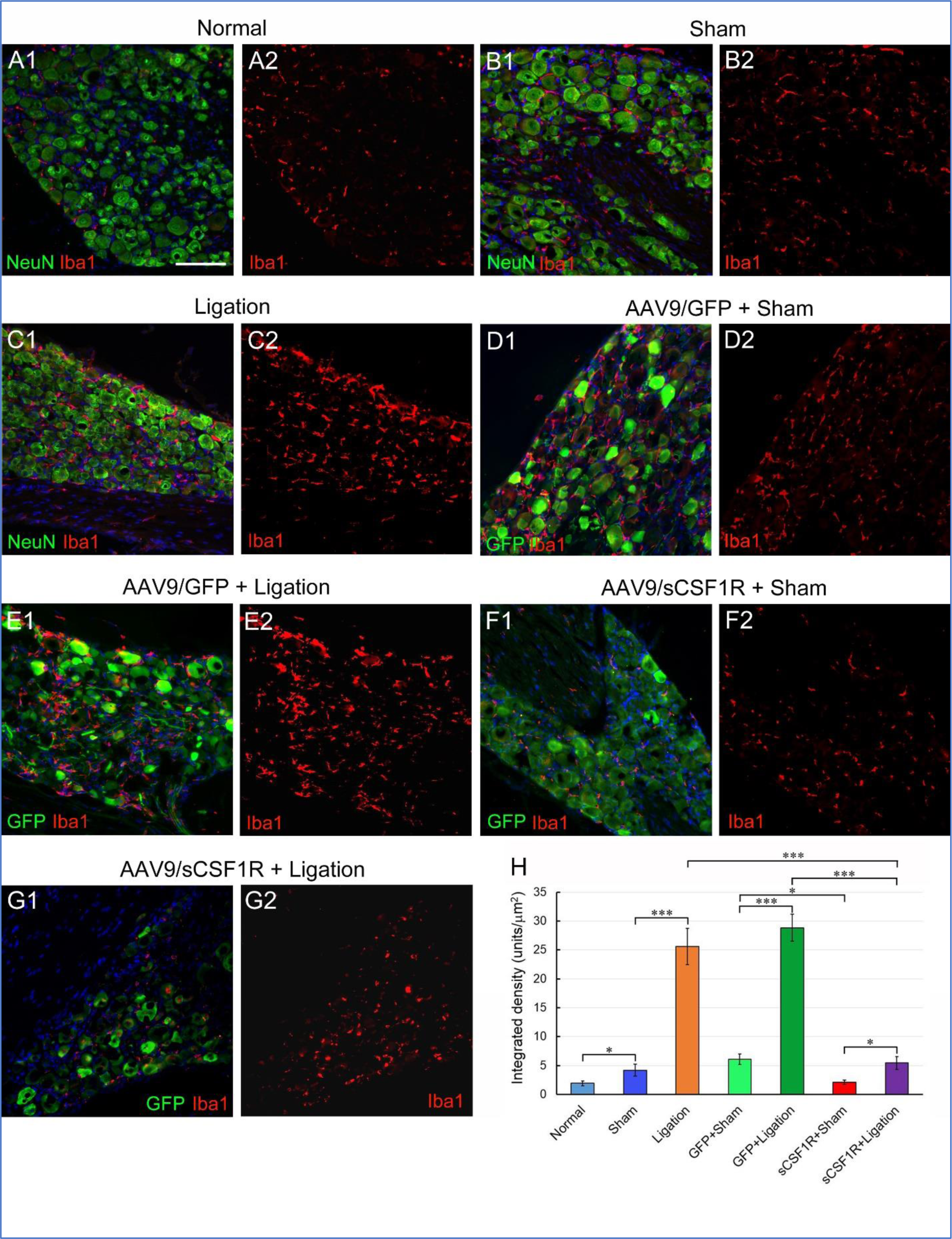
Over-expression of sCSF1R suppresses the elevation in macrophage densities in DRG after partial sciatic nerve ligation. Photomicrographs shown in this figure were taken from the left L4 DRG. 1, merged images; 2, red-channel only images. A1-A2 Normal. B1-B2 Sham-operated. C1-C2 Partial sciatic nerve ligation. D1-D2 AAV9/GFP + sham operation. E1-E2 AAV9/GFP + ligation. F1-F2 AAV9/sCSF1R + sham operation. G1-G2 AAV9/sCSF1R + ligation. H Quantification of the intensities of Iba1 immunoreactivity in DRG. n = 6 per group. Data information: Scale bar, 100 μm in (A1), also applies to the other photomicrographs. Data are presented as mean ± SEM. \**p* < 0.05; \*\*\**p* < 0.001. Statistic analysis was performed using one-way ANOVA with post doc Tukey test.

### Over-expression of sCSF1R suppresses macrophage accumulation of in sciatic nerve after partial sciatic nerve ligation

As sCSF1R expressed by DRG neurons would also be released into sciatic nerves, we examined the macrophages in the sciatic nerves. The areas chosen for Iba1-ir quantification were 2 mm proximal to the ligation sites or the corresponding locations from mice in the normal and sham surgery groups. A small number of macrophages were present in normal sciatic nerves (Fig 6A). Although sham surgery did not induce any significant change in macrophage density (Fig 6B and H), partial nerve ligation induced a significant increase in macrophage density (Fig 6C and H).

**Figure 6.**
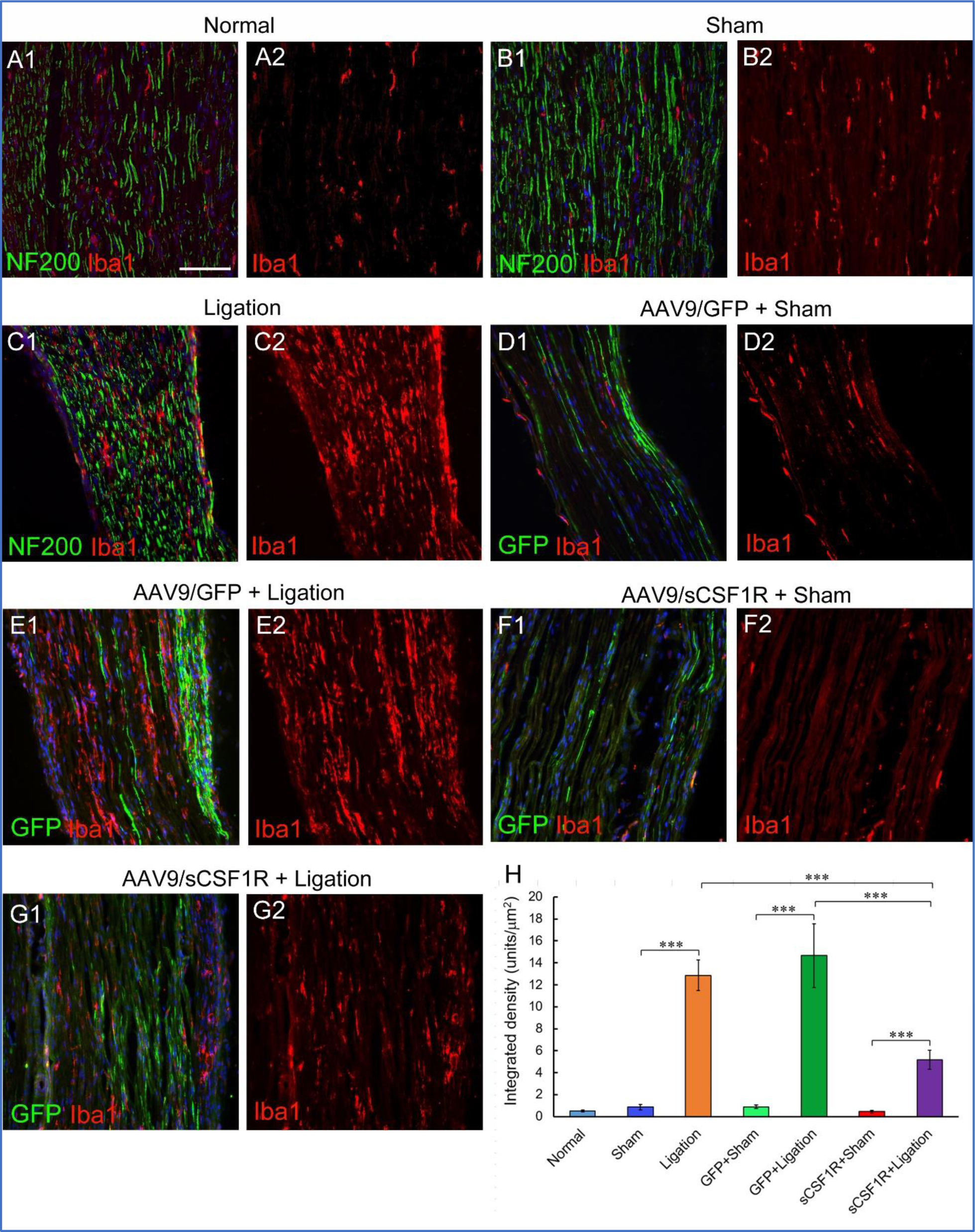
Over-expression of sCSF1R suppresses the elevation of macrophage densities in the sciatic nerve after partial sciatic nerve ligation. Photomicrographs shown in this figure were taken from sections about 2 mm proximal to the ligation site or the corresponding location in the mice of normal and sham surgery groups. 1, merge images; 2, red-channel only images. A1-A2 Normal. B1-B2 Sham surgery. C1-C2 Partial sciatic nerve ligation. D1-D2 AAV9/GFP + sham surgery. E1-E2 AAV9/GFP + ligation. F1-F2 AAV9/sCSF1R + sham surgery. G1-G2 AAV9/sCSF1R + ligation. H Quantification of the intensities of Iba1 immunoreactivities in the sciatic nerve. n = 6 per group. Data information: Scale bar, 100 μm in (A1), also applies to the other photomicrographs. Data are presented as mean ± SEM. \*\*\**p* < 0.001. Statistic analysis was performed using one-way ANOVA with post doc Tukey test.

As expected, the sham surgery on AAV9/GFP-injected mice did not lead to a significant increase in macrophage density (Fig 6D and H). However, partial nerve ligation on AAV9/GFP-injected mice induced a very significant increase in macrophage density (Fig 6E and H). Sham surgery on AAV9/sCSF1R injected mice did not significantly increase the macrophage density compared with the normal and the sham surgery groups (Fig 6F and H). Although nerve ligation on AAV9/sCSF1R-injected mice increased the macrophage density significantly compared with the sham-operated mice injected with AAV9/sCSF1R, the density was significantly lower than both the partial nerve ligation group and the AAV9/GFP plus nerve ligation group (Fig 6G and H). The results indicate that over-expression of sCSF1R in DRG neurons can significantly suppress the accumulation of macrophages in sciatic nerves induced by partial nerve ligation.

### sCSF1R attenuates mechanical allodynia induced by partial sciatic nerve ligation

Development of mechanical allodynia was assessed using the Von Frey hair test. Partial sciatic nerve ligation induced a significant decrease in paw withdrawal thresholds over the experimental 14-day period (Fig 7A), indicating the development of mechanical allodynia. Sham surgery involving the exposure of sciatic nerves was sufficient to induce a moderately decreased paw withdrawal threshold (Fig 7A). Sham surgery on both AAV9/GFP- and AAV9/sCSF1R-injected mice led to reduced paw withdrawal thresholds, however, they were not statistically significant from that of the sham surgery-only group (Fig 7A). AAV9/GFP injection followed by partial sciatic nerve ligation significantly reduced the paw withdrawal threshold, to a non-significant lower level than that of the partial nerve ligation-only group (Fig. 7A). AAV9/sCSF1R injection followed by partial nerve ligation also reduced the paw withdrawal threshold (Fig. 7A), but the extent of the reduction was much less in comparison with the AAV9/GFP plus partial nerve ligation group (Fig 7B). These results indicate that over-expression of sCSF1R can alleviate mechanical allodynia induced by partial nerve ligation.

**Figure 7.**
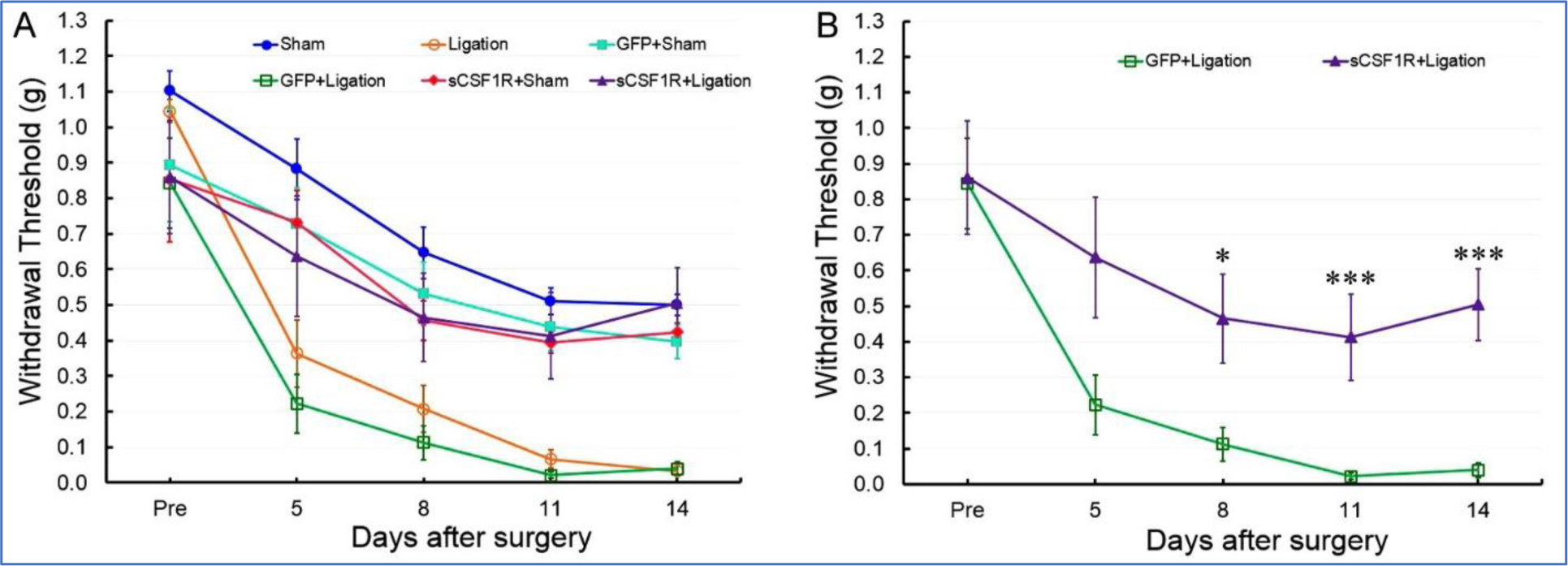
sCSF1R attenuates mechanical allodynia induced by partial sciatic nerve ligation. A Withdrawal thresholds of the ipsilateral hindlimbs when stimulated with von Frey filaments for all six experimental groups (results of statistical analysis are not shown). B To show the results of the statistical analysis clearly, only the partial nerve ligated groups treated with the AAV9/GFP and AAV9/sCSF1R are presented. Data information: Data are presented as mean ± SEM. n = 6 per group. \**p* < 0.05, \*\*\**p* < 0.001, compared with the same time point between the two groups. Statistic analysis was performed using two-way ANOVA with post doc Tukey test.

## Discussion

Prevention of proliferation and activation of microglia in the spinal cord and accumulation of macrophages in DRG and injured nerves has been implicated as an effective approach in the alleviation of neuropathic pain caused by peripheral nerve injuries. Although several CSF1R inhibitors have been shown to deplete microglia in the CNS, it is not clinically plausible to deplete or significantly reduce the number of microglia, monocytes, and resident macrophages in all tissues in order to alleviate neuropathic pain, as doing so would significantly weaken the body’s immune system. Therefore, localized and long-term delivery of therapeutic molecules that can suppress the proliferation and activation of microglia and accumulation of macrophages in the affected tissues would be the desired approach for treating injury-induced acute and chronic neuropathic pain. The results from the current study demonstrate that our approach of viral vector-mediated expression of a non-functional sCSF1R in DRG and the spinal cord addresses both the spatial and temporal issues for a feasible treatment of neuropathic pain.

One crucial issue that determines the feasibility of our approach is whether sufficient sCSF1R can be secreted *in vivo* to neutralize CSF1. CSF1R is also one of the receptors for interleukin 34 (IL-34) which plays an important role in the development and maintenance of microglia and Langerhans cells in the epidermis (Greter *et al*, 2012; Wang *et al*, 2012). IL-34 is highly expressed in the mouse CNS as we have previously reported (Gushchina *et al*., 2018). Therefore, IL-34 would compete with CSF1 for binding to sCSF1R as the two ligands have similar, although different, binding sites on the CSF1R (Nakamichi *et al*, 2013). To efficiently target the sCSF1R to the constitutive secretory pathway for high-level secretion, we fused the preproNGF signal peptide to sCSF1R cDNA. Previously we have successfully targeted a soluble Nogo receptor to the constitutive secretory pathway using the full-length NGF molecule (Zhang *et al*, 2013). Using only the signal peptide of NGF we were able to achieve the same goal with the sCSF1R in this study. Immunoblotting results confirmed that a high level of sCSF1R was present in the conditioned medium of HEK293 cells transfected with the plasmid carrying the sCSF1R construct. Detection of sCSF1R in the cerebrospinal fluid also indicates efficient secretion of sCSF1R *in vivo*.

We found that intrathecal injection of AAV9 into the lumbar spine predominantly transduced the lumbar and sacral DRG, which is desirable for this study as the mouse sciatic nerve mainly derives from the L3-5 spinal nerve roots. The transduction efficiency of AAV9/GFP and the distribution of transduced cells and axons are similar to those reported by Schuster and colleagues using the same strain of mice and a similar dose of AAV9/GFP (Schuster *et al*, 2014). Having a high transduction efficiency of DRG neurons would enable sCSF1R to be expressed in the same sensory neurons with increased CSF1 expression. CSF1 secreted into the spinal cord by the central axons of DRG neurons, inside DRG, and in sciatic nerves would bind to the high level of sCSF1R rather than activating the native CSF1R on microglia and macrophages, leading to reduced microgliosis and inflammation. An unexpected finding was that over-expression of sCSF1R in DRG neurons suppressed the elevation of CSF1 expression in DRG neurons induced by nerve ligation. One potential reason is that both CSF1 and CSF1R were targeted to the same secretory vesicles where they were bound to each other, resulting in the degradation of CSF1 in the cells. On the other hand, the CSF1 level in motor neurons in the ventral horns was also reduced. As most of these motor neurons were not transduced by AAV9/sCSF1R, it indicates that the effect of sCSF1R in suppressing CSF1 elevation inside the motor neurons was due to reduced inflammation in sciatic nerves.

As expected, partial sciatic nerve ligation in our study led to microgliosis in the dorsal and ventral horns of the lumbar spinal cord, as previously reported by other groups (Colburn *et al*., 1999; Lee *et al*., 2018; Okubo *et al*., 2016; Zhang *et al*, 2008). Over-expression of sCSF1R significantly decreased the extent of microgliosis in both dorsal and ventral horns, implicating that sCSF1R can reduce the activation of native CSF1R on microglia. The microglia densities in both dorsal and ventral horns were still higher than those of untreated mice or mice subjected to sham surgery, indicating that the sCSF1R level in this study may not be high enough to completely prevent microgliosis. Higher doses of AAV9/sCSF1R may be tested in a future study, but complete blockage of CSF1R is not our intention as our goal is to control the proliferation and activation of microglia to alleviate neuropathic pain, not depletion of microglia as they have many physiological roles.

Although recent studies on the role of CSF1 in the development of neuropathic pain has been focused on microgliosis in the spinal cord, the involvement of macrophages in the development of neuropathic pain has also been investigated in recent years (Ristoiu, 2013). An increase in macrophage density in DRG following traumatic peripheral nerve injury was reported nearly three decades ago (Lu & Richardson, 1993). Other types of nerve injuries, such as metabolic diseases, neurotoxins, and viral infection, can also result in increased macrophage densities in DRG. Since macrophages respond rapidly to peripheral nerve injuries, they are considered the initiators of neuropathic pain (Peng *et al*, 2016; Yu *et al*., 2020). Hematogenous macrophages can infiltrate the DRG after nerve injury and both hematogenous and resident macrophages proliferate in the DRG, in which the increased expression and secretion of CSF1 may play an important role. CSF1 was shown to promote the proliferation of macrophages and also act as a chemotactic factor to recruit monocytes from blood (Pixley, 2012). It should be noted that monocyte chemoattractant protein-1 was reported to be induced in Schwann cells in a few hours after nerve injury and is considered the main chemotactic factor for recruiting monocytes into injured nerves (Subang & Richardson, 2001). The macrophages were observed to be distributed within the DRG or surround the neurons (Lu & Richardson, 1993; Vega-Avelaira *et al*, 2009). A potential suggestion for the macrophages surrounding the neurons might be due to the action of the secreted CSF1 from the corresponding neurons. In our previous study on the EAE model of multiple sclerosis, we found that activated microglia surrounded those motor neurons that expressed a high level of CSF1 (Gushchina *et al*., 2018). In this study, we also observed that activated microglia surrounded motor neurons in ventral horns and macrophages surrounded sensory neurons in DRG in the mice injected with AAV9/GFP and with partial sciatic nerve ligation (Fig EV2H and EV3H). Secretion of inflammatory factors, such as TNF-α, interleukin-1β, and interleukin-6, from macrophages close to the sensory neurons would sensitize the neurons to stimulations and result in neuropathic pain (Ellis & Bennett, 2013; Thacker *et al*, 2007). In the current study, sCSF1R expressed by DRG neurons effectively suppressed the increase of macrophage numbers in DRG induced by nerve ligation. Neutralizing CSF1 in DRG would reduce the penetration and proliferation of macrophages, hence reduce inflammation in DRG, which would, in turn, suppress the elevation of CSF1 expression in DRG neurons resulting in a reduced level of CSF1 in the dorsal horn and diminished microgliosis.

In addition to those in the DRG, macrophages in the injured nerves have also been reported to contribute to the development of neuropathic pain. Following traumatic nerve injury, resident macrophages and Schwann cells initiate inflammatory responses and recruit circulating leukocytes to the site of injury. In the early stage of physical peripheral nerve injury, the roles of macrophages, together with dedifferentiated Schwann cells, are to phagocytize cell debris and promote axonal regeneration. However, the accumulated macrophages in the injured nerves can also release inflammatory factors and contribute to the development of neuropathic pain (Kiguchi *et al*, 2017). Reducing the cytokine and chemokine releasing-macrophages in the injured nerves was shown to hamper the development of neuropathic pain in a partial sciatic nerve ligation model (Echeverry *et al*, 2013). Our data show that sCSF1R expressed in DRG neurons can significantly reduce the number of macrophages in the sciatic nerves proximal to the ligation sites, which would contribute to the alleviation of neuropathic pain, together with reduced macrophage density in DRG and reduced microgliosis in the dorsal horns.

In this study, the paw withdrawal thresholds were generally proportional to the extent of microgliosis in dorsal horns and densities of macrophages in DRG and sciatic nerves, further confirming the roles played by microglia and macrophages in the development of mechanical allodynia reported in other studies. Animals treated with AAV9/sCSF1R had a significantly higher threshold than those treated with AAV9/GFP, proving our postulation on the efficacy of sCSF1R in alleviating neuropathic pain. However, there is a certain discrepancy between the behavioural data and histological data. In AAV9/sCSF1R-injected mice, the microglia density in the dorsal horn and the macrophage densities in DRG and sciatic nerves in the nerve ligation group were significantly higher than those in the sham surgery group. However, there was no statistical significance in paw withdrawal thresholds between these two groups. One possible reason is that sham surgery as a type of trauma itself caused increased sensitivity to mechanical stimulation, which reduced the potential to detect any difference between the two AAV9/sCSF1R-treated groups. Sham surgery involves the injury to the tissues overlying the sciatic nerves, which could lead to the accumulation of inflammatory cells and release of inflammatory factors, causing irritation to sciatic nerves. A future study on a chronic nerve ligation model would shed light on the reason for such a discrepancy, as the wound caused by surgery would have healed and the local inflammation in the soft tissues around the nerves subsided.

In this study, we have proven the efficacy of sCSF1R in alleviating acute neuropathic pain. However, the final goal is to develop sCSF1R into a treatment for patients with chronic neuropathic pain. It has been reported that microglia and monocytes synergistically promote the transition from acute to chronic pain after nerve injury (Peng *et al*., 2016). In another study on rats it was suggested that microglia are required for maintaining long-term (>3 months) neuropathic pain after partial sciatic nerve ligation (Echeverry *et al*., 2017). As AAV mediated gene expression in neurons can last for years, AAV9/sCSF1R treatment should be able to suppress the microgliosis and the accumulation of macrophages over a long period and effectively alleviate neuropathic pain.

Mounting evidence indicates that microglia also participate in the pathogenesis of several neurological disorders, especially neurodegenerative diseases (Hickman *et al*, 2018). CSF1 and CSF1R were reported to be involved in tau-induced neurodegeneration (Mancuso *et al*, 2019), Alzheimer’s disease (Murphy *et al*, 2000), amyotrophic lateral sclerosis (Gowing *et al*, 2009), Parkinson’s disease (Oh *et al*, 2020), spinocerebellar ataxia type 1 (Qu *et al*, 2017), Huntington’s disease (Crapser *et al*, 2020), and multiple sclerosis (Gushchina *et al*., 2018; Nissen *et al*, 2018). Over-expression of sCSF1R in the CNS could be employed to prevent the activation of microglia, hence slowing down the progression of these neurodegenerative disorders, therefore, the gene therapy of using sCSF1R could have broader clinical applications.

In conclusion, this study shows that AAV9-mediated expression of sCSF1R in DRG and the spinal cord can successfully reduce microgliosis in the spinal cord and suppress the accumulation of macrophages in DRG and injured nerves after partial sciatic nerve ligation. Such an approach targets the processes for both the central and peripheral sensitization of the sensory nerves, leading to the alleviation of neuropathic pain. Overall, the findings from this study may form the base for the future development of sCSF1R into gene therapy for treating neuropathic pain, and potentially, expanding to neurodegenerative diseases.

## Materials and methods

### Construction of expression cassette for sCSF1R

A fusion molecular construct was made to produce a vector that can efficiently express and secrete sCSF1R as well as express a reporter for easy identification of the transduced cells. The signal peptide of human prepro-nerve growth factor (NGFsp) was fused to the N-terminal of the CSF1-binding domain of CSF1R. NGFsp functions to direct sCSF1R proteins into the constitutive secretory pathway, so they can be secreted continuously. An HA-tag was attached to the C-terminal of the CSF1R extracellular CSF1-binding domain for the detection of sCSF1R by immunoblotting and immunoprecipitation. GFP is linked to the sCSF1R construct via a 2A peptide (from foot-and-mouth disease virus) at its N-terminal, which enables ribosome skipping at the 2A site that would lead to translation of sCSF1R and GFP separately. In this way, GFP remains in the transduced cells as a reporter while sCSF1R is secreted constantly. The NGFsp-sCSF1R-HA-2A-GFP fusion construct was generated using PCR and overlap extension. The mouse CSF1R cDNA (IMAGE ID: 30436119) was purchased from Source Bioscience. The DNA sequence encoding the extracellular domain of CSF1R (aa 1-506) was obtained by PCR with a 5’-primer containing the NGFsp sequence and the 3’-primer containing the HA sequence (Table 1). The HA-2A DNA fragment was obtained by amplifying two annealed primers: one with the 5’-HA sequence and the overlapping 2A sequence, the other with a complementary overlapping 2A sequence (Table 1). An overlap extension reaction was performed to generate the NGFsp-sCSF1R-HA-2A DNA construct, which was subcloned into pEGFP-N1 to form pNGFsp-sCSF1-HA-2A-GFP. The pAAV/CAG_GFP (Plasmid ID 37825) was purchased from Addgene, which has a hybrid chicken β-actin promoter (CAG) to drive the expression of GFP. The NGFsp-sCSF1R-HA-2A-GFP construct was subcloned into pAAV/CAG_GFP to generate pAAV/CAG_NGFsp-sCSF1R-HA-2A-GFP, which is abbreviated to pAAV/sCSF1R. The pAAV/CAG_GFP is abbreviated to pAAV/GFP. The molecular construct of the expression cassette is shown in Fig 1A.

**Table 1.**
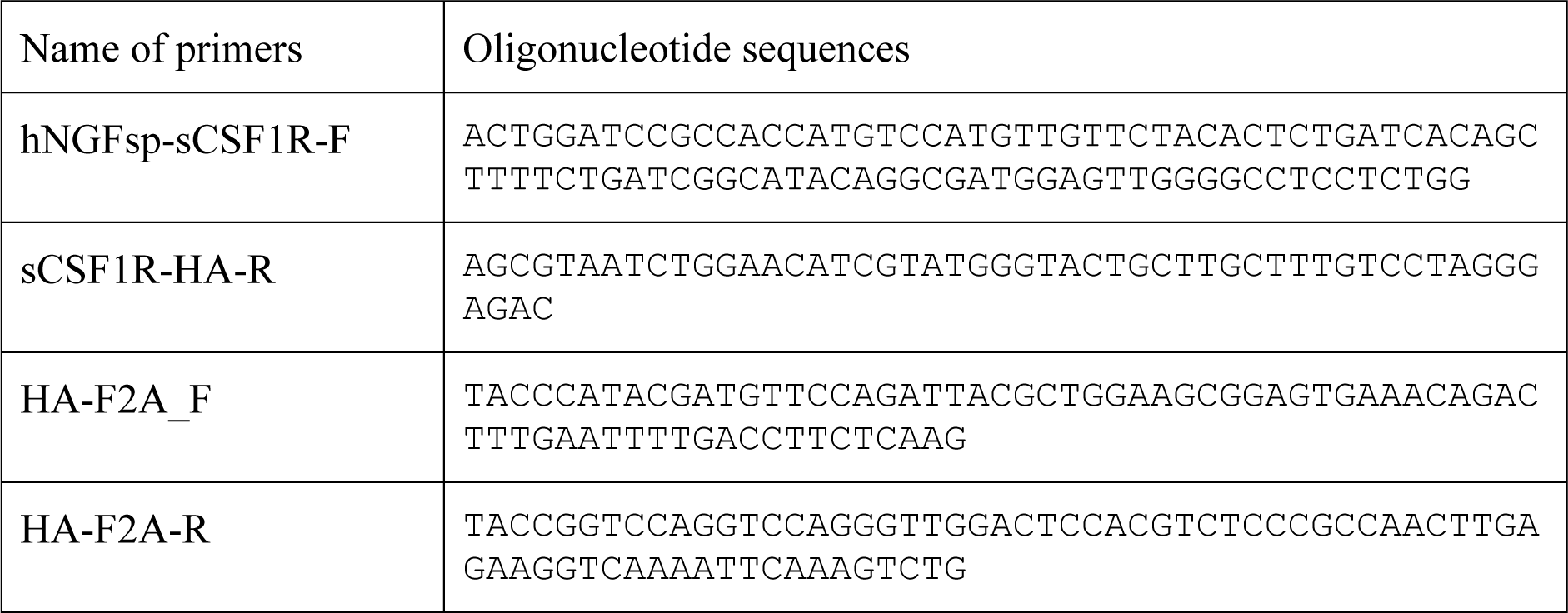
Sequences of PCR primers used to generate the hNGFsp-sCSF1R-HA-2A-GFP fusion construct

### Production of adeno-associated viral vectors

AAV9/GFP and AAV9/sCSF1R were produced mainly based on the protocols by Mulcahy and colleagues (Mulcahy *et al*, 2015). HEK293T cells were cultured in DMEM-GlutaMAX (Thermo Fisher Scientific) with 10% foetal bovine serum and penicillin (100 units/ml)/streptomycin (100 µg/ml). Cells were transfected with pAAV/GFP or pAAV/sCSF1R-GFP, pHelper, and pAAV9-RC (Gene Therapy Program, University of Pennsylvania) using linear polyethylenimine (PEI, MW. 25,000, Polyscience) with a PEI:DNA ratio of 4:1 in weight. Cells were harvested 72 h post transfection. Viral particles were released from cell pellets by four freeze-thawing cycles and purified by iodixanol gradient ultracentrifugation, followed by desalting and concentration using Amicon Ultra-15 100K Centrifugal Filter Units (Merck). Viruses were aliquoted and stored at -80°C. Titration of the viral vectors was performed by qPCR using a KAPA SYBR® FAST qPCR Master Mix Kit (Kapa Biosystems). The titre was 7.9 × 10^13^ GC/ml for AAV9/GFP, and 5.0 × 10^13^ GC/ml for AAV9/sCSF1R.

### Immunocytochemistry

To detect the expression of sCSF1R *in vitro*, HEK293T cells were cultured on 13 mm round coverslips in 35 mm dishes and transfected with pAAV/sCSF1R using Lipofectamine 2000 (Thermo Fisher Scientific). One day after transfection, cells were fixed with 4% paraformaldehyde and washed with phosphate-buffered saline (PBS). Cells on coverslips were incubated with Blocking Solution (10% goat serum, 0.2% Tween-20 in PBS) for 1 h and then incubated with a rabbit antibody against the extracellular domain of CSF1R (Merck, 06-174,) or a mouse anti-HA tag antibody (BioLegend, 901501) for 3 h at room temperature. After washing in PBS, cells were incubated with an Alexa Fluor-568 conjugated goat anti-rabbit or anti-mouse antibody (Thermo Fisher Scientific) for 1 h. DAPI-Fluoromont-G (SouthernBiotech) was applied to the coverslip to stain the nuclei and mount the coverslip onto a slide.

### Immunoblotting, immunoprecipitation, and co-immunoprecipitation

To investigate the secretion of sCSF1R into the conditioned medium, HEK293T cells were cultured in Animal-Free PeproGrow AF-CHO Media (PeproTech) in a 10 cm dish and transfected with pAAV/sCSF1R using Lipofectamine 2000. Two days after transfection, media were harvested and concentrated by 20 times using Amicon Ultra-15 Centrifugal Filter Units (Ultracel-10K, Merck).

Conditioned medium (2 µl per lane) was mixed with reducing loading buffer, heated at 70°C for 10 min, and loaded into a 4-12% SDS-PAGE precast gel (Abcam). Proteins were transferred on to PVDF membrane, which was incubated in blocking buffer for 1 h (10% skimmed milk in TBST buffer (Tris-buffered saline + 0.3% Tween 20)). The membrane was then incubated with rabbit anti-CSF1R antibody (1:1500, Merck), or rabbit anti-GFP (1:40,000, Abcam, ab290), or mouse anti-HA (1:5000, BioLegend) overnight, followed by incubation with donkey anti-rabbit IgG-HRP (1:10,000, GE Healthcare) or sheep anti-mouse IgG-HRP (1:10,000, GE Healthcare) for 2 h. Lumigen ECL Ultra (Lumigen) was used for signal generation and the ECL signal was recorded with ChemiDoc™ XRS+ System (BioRad).

Immunoprecipitation was performed to confirm that the protein bands identified in immunoblot were indeed sCSF1R. Thirty μl of PureProteome™ Protein A/G Mix Magnetic Beads (Merck) was washed in TBST and then incubated with 5 μl anti-HA antibody for 60 min at room temperature. Free antibodies were removed by washing three times. Ten μl of sCSF1R conditioned medium was mixed with the beads and rotated at 4°C overnight. Beads were washed with TBST twice and 30 μl reducing sample buffer was added and heated at 70°C for 10 min. Fifteen μl of eluent from the beads was loaded in the gel. The rest of the procedure was the same as described above for immunoblotting.

Co-immunoprecipitation was performed to verify that sCSF1R could bind CSF1. After formatting the Protein A/G beads-anti-HA antibody-sCSF1R complex, 2 or 6 ng of recombinant murine CSF1 (mCSF1) (PeproTech) was added and incubated for 1 h at room temperature. After removal of the unbound mCSF1, the beads were washed with TBST twice rapidly to limit the detachment of the bound mCSF1 from sCSF1R. For control, beads were incubated with 6 ng of mCSF1 and washed twice rapidly. Twenty μl of Sample Buffer was added to each sample and heated at 70°C for 10 min. All 20 μl of eluent from the beads was loaded into a 4-12% SDS-PAGE precast gel. mCSF1 (1 ng) was also loaded into the gel, serving as the positive control.

The gel-running and blotting procedures were the same as above. For detection of mCSF1 blotted on to the membrane, the membrane was incubated with anti-mCSF1 antibody (1:1000, PeproTech, 500-P62G) at 4°C overnight. Donkey anti-goat IgG-HRP (1:10,000, Santa Cruz Biotechnology) was applied to the membrane for 2 h at room temperature.

### Intrathecal injection of viral vectors

Experiments were performed in accordance with the United Kingdom Animals (Scientific Procedures) Act 1986 and approved by the UK Home Office, following local ethical review of procedures. Adult male mice (23 - 28 g) were randomly allocated into three treatment groups: no virus injection, AAV9/GFP injection, and AAV9/sCSF1R injection. They were then coded and the operators were blind to the codes. The surgery was performed similarly as previously described (Barritt *et al*, 2006; Peluffo *et al*, 2013). Under isoflurane anaesthesia, a laminectomy was performed at the T11 spinal vertebra. After making a small incision in the dura, the tip of a 20 mm length of flexible micro-renathane tubing (0.25 mm OD × 0.13 mm ID, Braintree Scientific) connected to a 20 mm Silastic catheter (0.94 mm ID × 0.51 mm ID, VWR) was inserted 4 mm subdurally for delivery of the viruses, so that the tip of the tubing was around the L3 spinal segment level. Five μl bolus of AAV9/sCSF1R or AAV9/GFP (2.5 × 10^11^ GC) was injected at the rate of 1 μl/min using a Hamilton syringe connected to a microinfusion pump. Thereafter, the muscle and skin incisions were sutured. Animals were monitored daily after surgery and provided with analgesia twice daily for two days. Animals were kept at 12 h light/dark cycle with water and food available *ad libitum*. Three randomly selected mice were injected with AAV9/sCSF1R for collecting fresh tissues and cerebrospinal fluid for immunoblotting to detect the expression levels of sCSF1R in the CNS.

### Partial sciatic nerve ligation

Two weeks after intrathecal injections, one group of AAV9/GFP (n = 6) and one group of AAV9/sCSF1R (n = 6) injected mice were randomly selected and anaesthetized with isoflurane. The sciatic nerve in the left thigh was exposed and approximately 1/3 to 1/2 of the nerve width was tied off by passing a needle attached to 5-0 VICRYL^®^ suture (Johnson & Johnson) through the distal sciatic nerve as was described by Seltzer and colleagues (Seltzer *et al*., 1990). Two other groups of AAV9/GFP and AAV9/sCSF1R (n = 6 each) injected mice received sham surgery with sciatic nerve exposure only to serve as controls for nerve ligation. Partial sciatic nerve ligation was performed on the fifth group of mice without viral vector injection for control. The sixth group of mice was subjected to sciatic nerve exposure only.

### Behavioural test

The noxious mechanical threshold of the hind paw was determined with the von Frey hairs (Touch Test^®^, North Coast Medical) applied onto the plantar surface of the hind paw similarly to previously described (Staniland *et al*, 2010). Unrestrained animals were acclimatized on top of a uniform metal grid surface within plastic cubicles (8 × 5 × 10 cm) for at least 60 min before testing. Various sizes of von Frey hairs, starting with the midrange 0.6 g filament, were applied to the plantar surface of the hind paw for 3 sec or until the paw was withdrawn in a reflex not associated with movement or grooming. Filaments were applied alternately to the left and right hind paws. A 50% withdrawal threshold was calculated using the ‘up-down’ method (Chaplan *et al*, 1994). This up-down method initially involves the identification of positive or negative responses with the 0.6 g von Frey hair. If there was a response, then the next lower force hair was applied and vice versa until a change in response was observed. Thereafter, four subsequent hairs were then assessed according to the up-down sequence. A 50% paw withdrawal value was calculated using the method described by Dixon (Dixon, 1980).

Animals were accommodated in our Biological Service Unit for a week after delivery. They were then acclimated to the testing device by being placed in them once every day for 20 min for 3 days. Formal tests were then carried out on day 6, 5, 2, and 1 prior to the intrathecal injections, then on days 5, 8, and 13 post-viral injections. After sciatic nerve ligation or sham surgery, they were tested on day 5, 8, 11, and 14 post-injury. For sham surgery and ligation-only groups, they were tested as the same as those groups with AAV9 injections.

### Tissue sample collection

The mice were killed by intraperitoneal injection of Lethobarb on the 15^th^ day after partial sciatic nerve ligation or sham surgery. To obtain the samples for immunoblotting, cerebrospinal fluid was collected from the cisterna magna using a micro-syringe, and the spinal cord and brain were removed and frozen on dry ice immediately. To obtain the samples for immunohistochemistry, the mice were perfused transcardially with PBS, followed by 4% paraformaldehyde. Sciatic nerves, DRG, spinal cords, and brains were removed and post-fixed in 4% paraformaldehyde overnight. They were then cryopreserved in 30% sucrose before being embedded in OCT Compound for sectioning.

### Immunohistochemistry

Cryostat sections (12 µm thick) of the fixed tissues were cut and either used immediately or stored at -20°C. For AAV9/GFP and AAV9/sCSF1R-GFP injected mice, the tissue sections were double-stained with a rabbit anti-GFP antibody (Abcam) or a sheep anti-GFP (BioRad, 4745-1051) and an antibody for microglia (anti-Iba1, Wako, 019-19741 or Novus, NB100-1028), or neurofilament (anti-NF200, Sigma-Aldrich, N0142), or neurons (anti-NeuN, Abcam, ab177487) (see Table 2 for detailed information on the primary antibodies used). The immunosignals were detected with two different fluorophore-conjugated secondary antibodies. Incubation was carried out in a humidified chamber at room temperature and washing was carried out for 3 × 10 min in TBST buffer (0.1% Tween-20). DAPI-Fluoromont-G was used for nucleus-staining and slide-sealing.

**Table 2.**
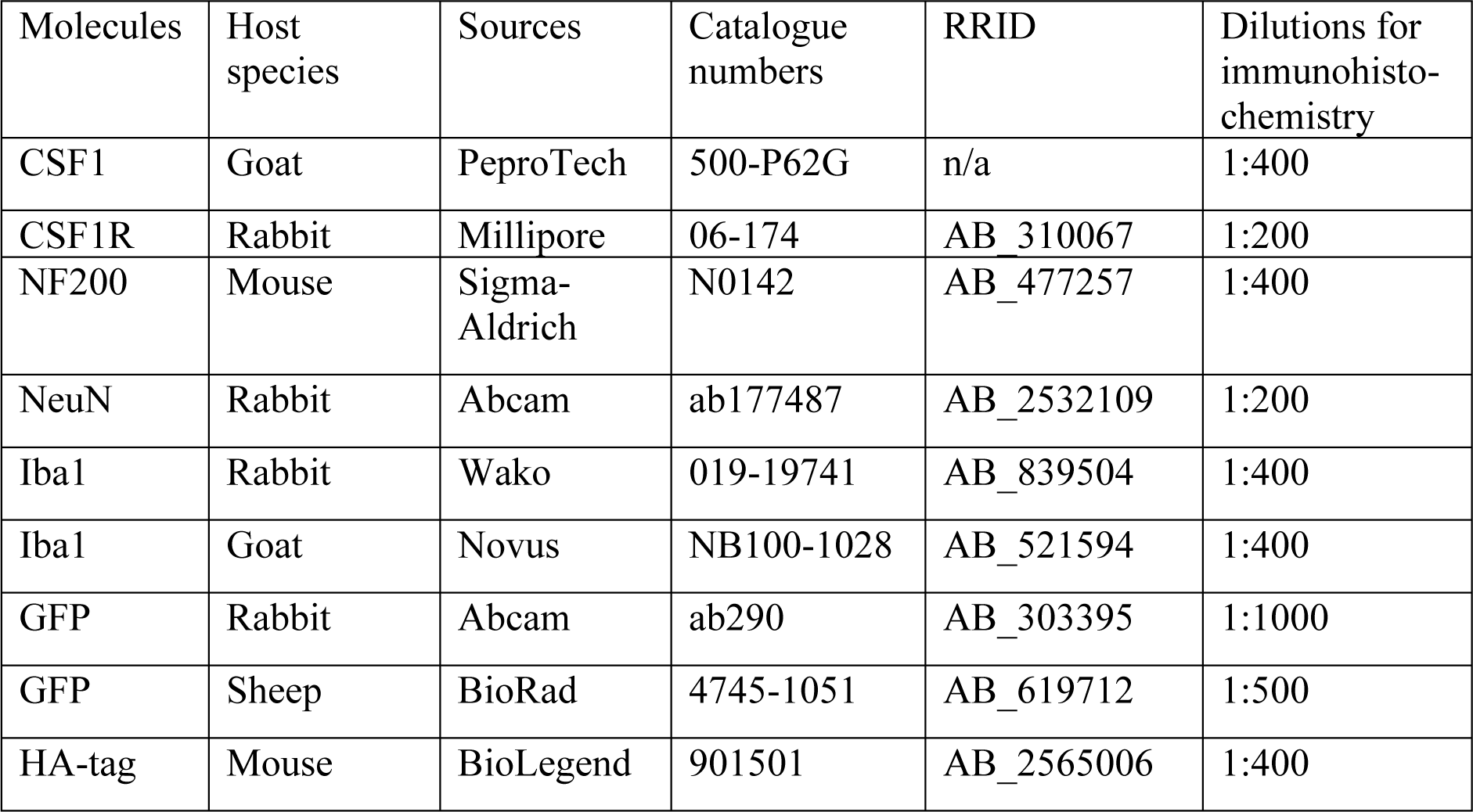
Primary antibodies used for immunohistochemistry and immunoblotting

For immunostaining of CSF1, sections were incubated with Blocking Solution 1 (10% horse serum, 0.3 M glycine, 0.3% Triton X-100 in PBS) at room temperature for 2 h. Goat anti-mCSF1 antibody (PeproTech) was applied and incubated at room temperature overnight. The sections were then incubated with polymer-HRP-conjugated horse anti-goat IgG antibody (ImmPRESS™ HRP Anti-Goat IgG Polymer Detection Kit, Vector Laboratories) for 30 min. Immunosignals were detected by incubating with ImmPACT™ DAB Substrate (Vector Laboratories).

To quantify the Iba1^+^ microglia in the lumbar spinal cords, one section from every four consecutive L4 spinal cord cross-sections was photographed with a 10× objective, and the integrated density of Iba1-ir in a frame of 330 × 450 µm in the dorsal horn on both ipsilateral and contralateral sides was measured using ImageJ (NIH). Four sections per animal were measured and the data were pooled.

To quantify the macrophages in DRG and sciatic nerves, ImageJ was used to measure the integrated densities of Iba1-ir. Images were acquired with a 20× objective. Due to the irregular shapes of DRG and sciatic nerve sections, the measuring areas were drawn manually to exclude the areas without tissues. Both the integrated density and the size of the measured area were recorded. The integrated density per µm^2^ was used for data analysis. Three sections with an interval of 36 µm between them from each animal were measured and averaged.

### Immunoblotting of tissue samples

Immunoblotting was performed to detect the relative levels of sCSF1R-HA-2A in the brain, spinal cord, and cerebrospinal fluid. Tissues were weighed and minced, and then tissue homogenization buffer (PBS containing 0.04% bovine serum albumin, Complete Protease inhibitors (Roche)) was added at 5 volumes of the tissue weight. Tissues were homogenized by sonication (30 W, 3 × 15 sec) using a sonicator (Vibra-Cell Ultrasonic Liquid Processor, Sonics & Materials). Homogenates were centrifuged at 10,000*g* for 20 min at 4°C and the supernatants were collected.

Ten μl of supernatant from the brain, spinal cord (cervical and thoracic segments, lumbar and sacral segments), 5 μl of cerebrospinal fluid, and 0.5 μl of conditioned medium from psCSF1R-transfected HEK293 cells were loaded into a 4 - 12% gradient gel. The rest of the procedure was the same as those described above. Anti-HA antibody (1:5000) was used to detect the sCSF1R-HA-2A on the blot.

### Statistical analysis

Data are presented as mean ± SEM. One-way ANOVA with post hoc Tukey test was used to compare the levels of Iba1-ir in the spinal cord, DRG and sciatic nerves. Two-way ANOVA with post hoc Tukey test was used to analyse the data for behavioural tests.

## Data availability

This study includes no data deposited in external repositories.

## Acknowledgements

We would like to thank Dongsheng Wu for performing the statistical analysis of the data and Lorna Bo for proofreading of the manuscript.

## Funding

The work was supported by a grant from Foresight Inc.

## Author contributions

XB supervised the entire project, designed the experiments, and wrote the manuscript. SG carried out most of the experiments and analysed the data. PKY performed the animal surgery. GAP, HS, JL, and ML participated in the immunohistochemistry.

## Conflict of interest

The authors declare no conflict of interests.

## The paper explained

### Problem

Millions of people worldwide suffer from debilitating chronic neuropathic pain and the current treatments are not so effective. Previous studies implicate that microglial proliferation and activation in the spinal cord and macrophage accumulation in the sensory ganglia and peripheral nerves play an important role in the development of neuropathic pain. Colony-stimulating factor-1 (CSF1) released by the sensory neurons was found to be responsible for microgliosis and macrophage accumulation after peripheral nerve injury. Therefore, blocking the CSF1 signalling pathway would be an effective approach to alleviating neuropathic pain.

### Results

We used an adeno-associated viral vector (AAV9) to over-express a non-functional soluble CSF1 receptor (sCSF1R) in mouse sensory ganglia and spinal cord, followed by partial sciatic nerve ligation to induce neuropathic pain. Pre-treatment with AAV9/sCSF1R significantly reduced the microglia densities in the spinal cord and macrophage densities in sensory ganglia and sciatic nerves compared to nerve-ligated mice pre-treated with the control vector AAV9/GFP. Behavioural tests showed that nerve-ligated mice pre-treated with AAV9/sCSF1R had a significantly higher pain threshold than the control group, indicating the alleviation of neuropathic pain.

### Impact

This is the first attempt to employ a gene therapy approach to block the CSF1 signalling pathway, which may represent a novel strategy in long-term alleviation of neuropathic pain. Moreover, given the importance of CSF1 and microglia/macrophages in a variety of neurodegenerative diseases such as Alzheimer’s, this approach may also be explored for the potential treatments of neurodegenerative and neuroinflammatory disorders.

---

**Figure EV1.**
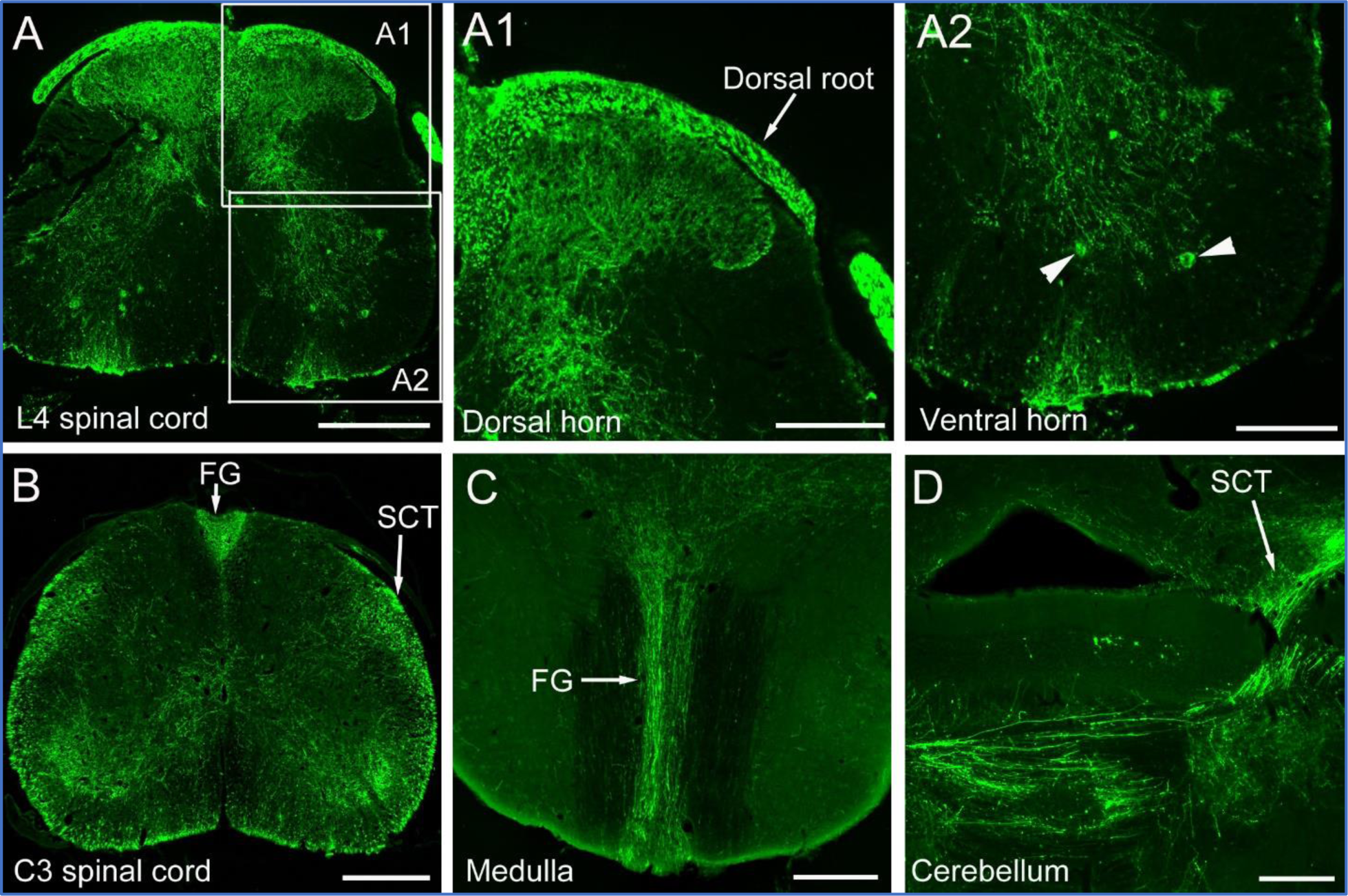
GFP expressing axons and neurons in spinal cord, medulla, and cerebellum in a naïve mouse injected with AAV9/GFP. The mouse was intrathecally injected with AAV9/GFP and sacrificed two weeks later. Sections were immunostained with an anti-GFP antibody. A A L4 spinal cord section. The two boxed areas are shown at higher magnification in (A1) and (A2). Arrowheads in (A2) point to two GFP-transduced motor neurons. B A C3 spinal cord section showing the GFP-expressing ascending sensory axons. The spinocerebellar tract (SCT) and fasciculus gracilis (FG) are indicated by arrows. C A horizontal section of medulla showing the GFP expressing axons of fasciculus gracilis. D A horizontal section of cerebellum showing GFP expressing axons of the spinocerebellar tract. Data information: Scale bar, 400 µm in (A - D); 200 µm in (A1) and (A2).

**Figure EV2.**
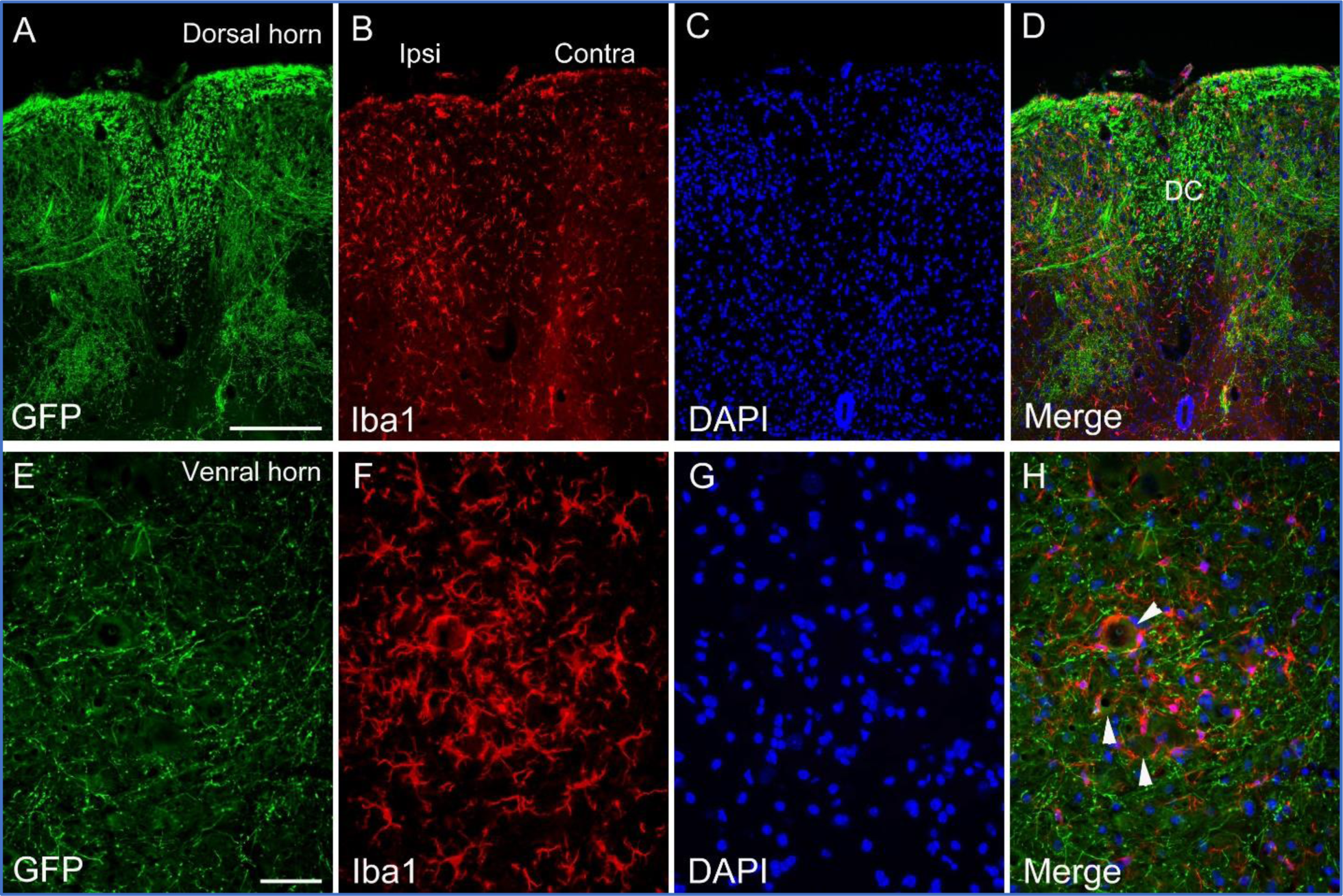
GFP expressing neurons and nerve fibres and microglia in the lumbar spinal cord in a mouse injected with AAV/GFP and received partial sciatic nerve ligation. The mouse was intrathecally injected with AAV9/GFP, followed by partial left sciatic nerve ligation two weeks later. The mouse was sacrificed two weeks after partial sciatic nerve ligation. Sections were immunostained for GFP and Iba1 (microglia). Nuclei were stained with DAPI. A–D An L4 spinal cord cross-section showing the GFP expressing sensory axons and Iba1-labelled microglia in the central region of the dorsal horns on both ipsilateral and contralateral sides to the partial sciatic nerve ligation. E-H An L4 spinal cord cross-section showing the GFP expressing sensory axons and Iba1-labelled microglia in the ventral horn on the ipsilateral side to the partial sciatic nerve ligation. Arrowheads in (H) point to the motor neurons surrounded by activated microglia. Data information: Ipsi: ipsilateral; contra: contralateral; DC: dorsal column. Scale bar, 200 µm in (A), also applies to (B–D); scale bar, 50 µm in (E), also applies to (F–H).

**Figure EV3.**
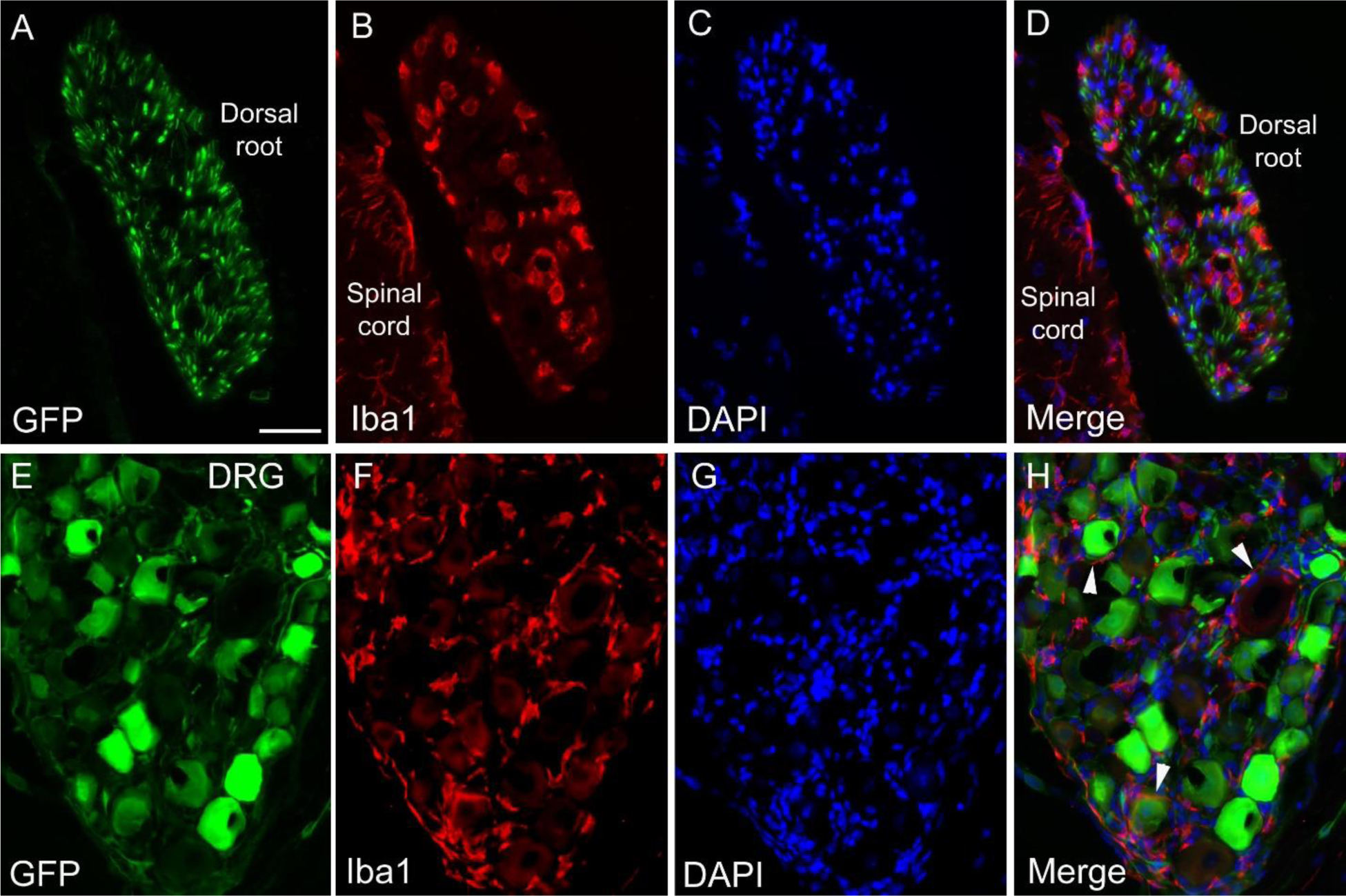
GFP expressing neurons and their axons and macrophages in dorsal root and dorsal root ganglion. The mouse was intrathecally injected with AAV9/GFP, followed by a partial left sciatic nerve ligation two weeks later. The mouse was sacrificed two weeks after sciatic nerve ligation. Sections were immunostained for GFP and Iba1 (macrophages and microglia). Nuclei were stained with DAPI. A–D GFP expressing sensory axons and Iba1-labelled macrophages in an L4 dorsal root. A small part of the spinal cord with Iba1-labelled microglia is also visible. E–H GFP expressing neurons and Iba1-labelled macrophages in an L4 dorsal root ganglion. Arrowheads in (H) point to neurons surrounded by macrophages. Data information: Scale bar, 50 µm in (A), also applies to (B–H).

